# Effectiveness and selectivity of a heroin conjugate vaccine to attenuate heroin, 6-acetylmorphine, and morphine antinociception in rats: Comparison with naltrexone

**DOI:** 10.1101/577494

**Authors:** Kathryn L. Schwienteck, Steven Blake, Paul T. Bremer, Justin L. Poklis, E. Andrew Townsend, S. Stevens Negus, Matthew L. Banks

## Abstract

**Background:** One emerging strategy to address the opioid crisis includes opioid-targeted immunopharmacotherapies. This study compared effectiveness of a heroin-tetanus toxoid (TT) conjugate vaccine to antagonize heroin, 6-acetylmorphine (6-AM), morphine, and fentanyl antinociception in rats.

**Methods:** Adult male and female Sprague Dawley rats received three doses of active or control vaccine at weeks 0, 2, and 4. Vaccine pharmacological selectivity was assessed by comparing opioid dose-effect curves in 50°C warm-water tail-withdrawal procedure before and after active or control heroin-TT vaccine. Route of administration [subcutaneous (SC) vs. intravenous [IV)] was also examined as a determinant of vaccine effectiveness. Continuous naltrexone treatment (0.0032-0.032 mg/kg/h) effects on heroin, 6-AM, and morphine antinociceptive potency was also determined as a benchmark for minimal vaccine effectiveness.

**Results:** The heroin-TT vaccine decreased potency of SC heroin (5-fold), IV heroin (3-fold), and IV 6-AM (3-fold) for several weeks without affecting IV morphine or SC and IV fentanyl potency. The control vaccine did not alter potency of any opioid. Naltrexone dose-dependently decreased antinociceptive potency of SC heroin, and treatment with 0.01 mg/kg/h naltrexone produced similar, approximate 8-fold decreases in potencies of SC and IV heroin, IV 6-AM, and IV morphine. The combination of naltrexone and active vaccine was more effective than naltrexone alone to antagonize SC heroin but not IV heroin.

**Conclusions:** The heroin-TT vaccine formulation examined is less effective, but more selective, than chronic naltrexone to attenuate heroin antinociception in rats. Furthermore, these results provide an empirical framework for future preclinical opioid vaccine research to benchmark effectiveness against naltrexone.

## 1. Introduction

Heroin abuse has been one major contributor to the current opioid crisis in the United States. In 2017, about 494,000 individuals aged 12 and older were determined to be current heroin users and approximately 652,000 people had an opioid-use disorder (OUD) with heroin as the primary abused drug (SAMHSA, 2018). In addition, the rate of heroin-related overdose deaths increased by almost 400% between 2010 and 2017 (Hedegaard et al., 2018). Current food and drug administration (FDA) approved treatments for OUD include the full and partial opioid agonists methadone and buprenorphine, respectively, and the opioid antagonist naltrexone. Although agonist-based therapies are effective, regulatory issues impede patient access (Jaffe and O’Keeffe, 2003). Recent estimates suggest only 21.5% of OUD patients received treatment in the previous year (Saloner and Karthikeyan, 2015). Patient compliance with naltrexone is notoriously poor even when naltrexone is combined with opiate abstinence incentives (Jarvis et al., 2019; Jarvis et al., 2018; Lee et al., 2018). Furthermore, some patients still relapse on current FDA-approved treatments (Barbosa-Leiker et al., 2018; Tkacz et al., 2012) highlighting the need for preclinical research to develop effective and readily accessible candidate OUD treatments.

Immunopharmacotherapies or “opioid vaccines” are one emerging treatment approach to address the ongoing opioid crisis (Baehr and Pravetoni, 2019; Banks et al., 2018; Volkow and Collins, 2017). An immunopharmacotherapy is defined as the use of highly selective antibodies, generated passively or actively, to sequester target drugs of abuse in the blood. Previous preclinical research supports the potential for a heroin-targeted vaccine to sequester heroin in the blood and attenuate centrally-mediated heroin effects (e.g. antinociceptive and reinforcing effects) (Anton and Leff, 2006; Bonese et al., 1974; Bremer et al., 2017; Matyas et al., 2014). Ethnographic reports indicate that heroin users may initiate heroin abuse via the intranasal route before eventually transitioning to intravenous (IV) heroin (Monico and Mitchell, 2018). Recently, vaccine effectiveness to attenuate brain oxycodone levels was greater when a large oxycodone dose was administered subcutaneously (SC) compared to IV in rats (Raleigh et al., 2018). Heroin is a prodrug rapidly metabolized to 6-acetylmorphine (6-AM) and morphine in the blood (Gottås et al., 2013; Inturrisi et al., 1984; Rook et al., 2006). SC heroin may allow for a great opportunity for hydrolysis to 6-AM than IV heroin administration, resulting in potentially greater heroin/6-AM antibody interactions. Consistent with this hypothesis, the heroin tetanus-toxoid (TT) conjugate vaccine evaluated in the present study displays a >10-fold affinity for 6-AM than heroin in mice and rhesus monkeys (Bremer et al., 2017). However, whether route of administration is an important determinant of heroin vaccine effectiveness remains to be determined.

The aim of the present study was to compare heroin-TT conjugate vaccine effectiveness between two routes of heroin administration (i.e. SC and IV) in male and female rats. A warm-water tail withdrawal procedure is one preclinical behavioral procedure to efficiently assess vaccine effectiveness and selectivity over time (Schlosburg et al., 2013; Stowe et al., 2011; Sulima et al., 2017). In addition, the present study also assessed heroin vaccine selectivity for the heroin metabolites 6-AM and morphine and the structurally dissimilar MOR (mu-opioid receptor) ligand fentanyl. For comparison, continuous naltrexone treatment effectiveness to attenuate SC and IV heroin antinociception was determined. In addition, naltrexone treatment selectivity to attenuate IV 6-AM, and IV morphine antinociception was also determined. Human laboratory studies and clinical trials suggest the minimally effective naltrexone dose produces an 8-10 fold potency shift in the heroin dose-effect function and maintains plasma naltrexone levels above 2 ng/ml (Bigelow et al., 2012; Comer et al., 2006; Sullivan et al., 2006). Thus, the present study also determined plasma naltrexone levels during continuous naltrexone treatments to correlate with heroin antinociception potency shifts.

## 2. Materials and Methods

### 2.1 Subjects

A total of 53 adult male (26) and female (27) Sprague Dawley rats (Envigo, Frederick, MD, USA) served as subjects. A total of 23 rats were surgically implanted with intravenous catheters (Townsend et al., 2019b). Estrous cycle was not monitored because recent guidelines and empirical data in our laboratory have suggested that when sex differences are present, they are often sufficiently robust for detection without reference to estrous cycle phase (Becker and Koob, 2016; Townsend et al., 2019b). All rats had unlimited access to rodent chow (Teklad LM-485 Mouse/Rat 7012, Envigo) and water in the home cage and were individually housed in a temperature- and humidity-controlled vivarium maintained on a 12-h light-dark cycle (lights on from 0600 to 1800). The vivarium was accredited by AAALAC International. Both experimental and enrichment protocols were approved by the Virginia Commonwealth University Institutional Animal Care and Use Committee in accordance with the National Institutes of Health Guidelines for the Care and Use of Laboratory Animals (Council, 2011).

### 2.2 Drugs

Heroin HCl, 6-AM, (−)-morphine sulfate, fentanyl HCl, and (−)-naltrexone HCl were supplied by the National Institute on Drug Abuse Drug Supply Program (Bethesda, MD). All drugs were dissolved in sterile water, except 6-AM which was dissolved in 1% lactic acid (Sigma Aldrich, St. Louis, MO). Drugs were administered SC or IV at an injection volume of 1-2 ml/kg and all drug doses were expressed as the salt forms listed except for 6-AM which was expressed as the base form. To convert the heroin dose reported as the HCl salt to free base heroin, multiply the HCl salt by (369.42/405.9) or 0.91. Drug solutions administered IV were filtered (0.22 μm sterile filter, Model # SLGV033RS, MilliporeSigma, Burlington, MA) before administration. For the continuous naltrexone experiments, naltrexone was delivered subcutaneously via osmotic pump.

### 2.3 Thermal nociception procedure

Rats were gently wrapped in an absorbent bench underpad (VWR, Radnor, PA) such that their tails hung freely. The bottom 5 cm of the tail was marked and immersed in water maintained at either 40 or 50°C in separate water baths (Thermo Precision 2833, Thermo Fisher Scientific, Waltham, MA, USA). Mercury thermometers were used to monitor water temperature. The latency for each rat to withdrawal its tail was measured by the same experimenter throughout the study using a hand-operated digital stopwatch with a time resolution of 1/100 s. If the rat did not remove its tail by 20 s, the experimenter removed the tail and a latency of 20 s was assigned. Baseline latencies at both 40 and 50°C were obtained in each daily test session before saline (vehicle) or drug administration. Cumulative-dose test sessions consisted of four to six cycles. Drugs were administered either SC or IV at the start of each cycle, and each drug dose increased the total cumulative dose by one-fourth or one-half log units. Saline was always administered first and was followed by drug administration. Tail-withdrawal latencies were redetermined 20 min later for SC experiments, 10 min later for IV heroin, 6-AM, and morphine experiments, and 1 min later for IV fentanyl experiments. All drugs were tested up to doses that produced maximal or near maximal antinociception (i.e., >80% maximum possible effect (%MPE)) or undesirable physiological effects such as respiratory depression that required naltrexone reversal. Drugs were tested no more than twice per week in an individual rat, and test sessions were separated by at least two days. Final sample sizes for each drug and treatment condition were generally three males and three females unless otherwise noted. Initially, dose-effect functions were determined for heroin (0.1-1mg/kg, SC; 0.032-1 mg/kg, IV), 6-AM (0.1-1 mg/kg, IV), morphine (0.32-10 mg/kg, IV), and fentanyl (0.001-0.032 mg/kg, IV), and each dose-effect function was determined once. Subsequently, two types of experiments were conducted.

### 2.4 Heroin vaccine experiments

First, a total of 18 rats (9 females, 9 males) were divided into three heroin vaccine experimental groups: 1) active vaccine/SC dosing, 2) active vaccine/IV dosing, and 3) control vaccine/IV dosing. Groups 1 and 2 compared whether route of heroin administration was a determinant of vaccine effectiveness. Groups 2 and 3 compared selectivity of the heroin vaccine for heroin and its metabolites 6-AM and morphine. Following initial baseline tail withdrawal experiments for heroin, 6-AM, morphine, and fentanyl, rats were vaccinated on a Friday at Week 0, 2 and 4. Testing continued throughout the vaccination period and was carried out for 8 weeks after initial vaccination in group 1, and 6-8 weeks after initial vaccination in groups 2 and 3.

The active heroin vaccine was composed of a heroin hapten that was conjugated to tetanus toxoid (TT) as previously described (Bremer et al., 2017) and solubilized in 50% glycerol/50% phosphate-buffered saline. Heroin copies were approximately 16-18 per protein. On a per-rat basis, 250 μg heroin-TT conjugate was mixed with 50 μg of phosphorothioate-modified CpG ODN 1826 (Eurofins Genomics, Louisville, KY) and 0.75 mg Alhydrogel adjuvant 2% (InvivoGen, San Diego, CA) for 30 min and then refrigerated for 24 hours prior to IP administration. Control vaccine consisted of TT (MassBiologics, Boston, MA) mixed with 50 μg of CpG ODN 1826 and 0.75 mg Alhydrogel. Every two weeks after initial vaccination, rats were sedated under isoflurane anesthesia and tail-vein blood was collected in a volume of approximately 0.5 mL into microtainer tubes without additive (BD, Franklin Lakes, NJ). Tubes were immediately centrifuged at 1000g for 10 min and the serum supernatant was transferred into a labeled storage tube and frozen at −80°C until analyzed for heroin mid-point titer levels as described previously (Bremer et al., 2017; Hwang et al., 2018a; Hwang et al., 2018c).

### 2.5 Chronic naltrexone treatment experiments

Heroin vaccine effectiveness was compared to chronic naltrexone administration to model aspects of clinically used depot naltrexone for OUD treatment. A total of 36 rats (18 males, 18 females) were divided into two experimental groups. Tail-withdrawal latencies following heroin, 6-AM, and morphine administration were determined using a 14-day experimental protocol. One group was tested with cumulative heroin (0.1-32 mg/kg, SC) before and on treatment days 7 and 14 following subcutaneous implantation of 14-day osmotic pumps (model 2ML2, 5 μL/h flow rate, Alzet, Cupretino, CA) filled with saline (n=6) or naltrexone (0.0032 mg/kg/h, n=6; 0.01 mg/kg/h, n=6; or 0.032 mg/kg/h, n=6). Another group was surgically implanted with intravenous jugular catheters and cumulative-heroin (0.032-10 mg/kg, IV) test sessions were conducted before and on days 7 and 14 following either saline (n=6) or naltrexone (0.01 mg/kg/h, n=6) pump implantation. After these initial heroin studies, pumps were aseptically removed on day 15 and a cumulative heroin dose-effect test session occurred 7 days later to determine whether heroin antinociceptive potency returned to pre-naltrexone levels. Subsequently, cumulative 6-AM (0.1-10 mg/kg, IV) and morphine (0.32-100 mg/kg, IV) dose-effect tests were determine before and on either day 7 or 14 following saline (n=5) and naltrexone (0.01 mg/kg/h, N=6) pump implantation. 6-AM and morphine were tested in a counterbalanced order such that half the rats received 6-AM on day 7 and the other half received morphine on day 7.

### 2.6 Naltrexone plasma sample collection and analysis

In addition to the naltrexone behavioral experiments, blood was collected during continuous 0.01 and 0.032 mg/kg/h naltrexone treatment on days 8 and 15. Rats were sedated under isoflurane anesthesia and approximately 0.5 mL of blood was collected from a tail vein into a microtainer tubes with lithium heparin (BD, Franklin Lakes, NJ). Tubes were immediately centrifuged at 1000g for 10 min and the plasma supernatant was transferred into a labeled storage tube and frozen at −80°C until analyzed. The plasma samples and quality control specimens prepared in plasma at low (3 ng/mL), mid (50 ng/mL) and high (350 ng/mL) concentrations of naltrexone were kept at −30°C until analyzed. A seven-point naltrexone calibration curve (1, 5, 10, 20, 100, 200 and 500 ng/mL), along with a drug free control and a negative control without internal standard (ISTD) in drug-free plasma was prepared. Naltrexone was extracted from the plasma using a liquid/liquid extraction. In brief, 100 ng/mL naltrexone-d_3_ and the ISTDs were added to 100 µL aliquots of plasma of each calibrator, control or specimen except the negative control. Then 0.5 mL of saturated carbonate/ bicarbonate buffer (1:1, pH 9.5) and 1.0 mL of chloroform:2-propanol (9:1) were added. Samples were mixed for 5 min and centrifuged at 2500 rpm for 5 min. The top aqueous layer was aspirated, and the organic layer was transferred to a clean test tube and evaporated to dryness at 40°C under a constant stream nitrogen. The samples were reconstituted with methanol and placed in auto-sampler vials for analysis. The ultra-performance liquid chromatography tandem mass spectrometer (UPLC-MS/MS) analysis was performed on a Sciex 6500 QTRAP system with an IonDrive Turbo V source for TurbolonSpray® (Sciex, Ontario, Canada) attached to a Shimadzu UPLC system (Kyoto, Japan) controlled by Analyst software (Sciex, Ontario, Canada). Chromatographic separation was performed using a Zorbax XDB-C18 4.6 x 75 mm, 3.5-micron column (Agilent Technologies, Santa Clara, CA). The mobile phase contained water:methanol (80:20, v/v) with 1 g/L ammonium formate and 0.1 % fomic acid and was delivered at a flow rate of 1 mL/min. The source temperature was set at 650°C, and curtain gas had a flow rate of 30 mL/min. The ionspray voltage was 5500 V, with the ion source gases 1 and 2 having flow rates of 60 mL/min. The declustering potential and collection energy were 86 eV and 39 eV, respectively. The quantification and qualifying transition ions were monitored in positive multiple reaction monitoring mode: Naltrexone 342> 267 and 342 > 282 and Naltrexone-d_3_ 345> 270 and 345 > 285. The total run time for the analytical method was 4 min. A linear regression of the peak area of ratios of the quantification transition ions and the ISTDs transition ions were used to construct the calibration curves.

### 2.7 Chronic naltrexone treatment after vaccination

Because of the chemical structure similarities between naltrexone, heroin, 6-AM, and morphine, an additional experiment was conducted in the active and control heroin-vaccinated rats to evaluate whether the heroin vaccine altered naltrexone effectiveness to antagonize heroin antinociception. For Group 1 rats (active heroin vaccine/SC administration), cumulative heroin dose-effect functions were conducted before and following implantation of 14-day osmotic naltrexone (0.01 mg/kg/h) pumps. Naltrexone pumps were implanted on a Friday 8 weeks post-initial heroin vaccine administration. Heroin dose-effect functions were determined twice in week 9, and data were averaged for presentation and analysis. For Group 2 rats (active heroin vaccine/ IV administration), cumulative heroin dose-effect functions were determined following implantation of 14-day osmotic naltrexone (0.01 mg/kg/h) pumps. Pumps were implanted at week 10, heroin was tested twice in week 11, and data were averaged for presentation and analysis.

### 2.8 Data Analysis

Raw tail-withdrawal latencies were converted to percent maximal possible effect (%MPE) using the equation: %MPE = [(Test latency – Saline latency) / (20 s – Saline latency)] X 100 where “test latency” was the latency from 50°C water after drug, and “saline latency” was the latency from 50°C after vehicle administration prior to drug administration. The ED_50_ value (and 95% confidence limits) of each dose-effect function was defined as the drug dose that produced 50%MPE and calculated using nonlinear regression using the [Agonist] vs. normalized response equation (GraphPad Prism, La Jolla, CA). Individual potency ratios were calculated by dividing the ED_50_ value during the test condition (vaccine or naltrexone treatment) by the baseline ED_50_ value and then averaged to yield group mean values. Group mean potency ratios were analyzed using a paired or unpaired Welch’s t-test, a RM one-way ANOVA, or two-way ANOVA, as appropriate. A Sidak post-hoc test followed all significant main effects. The criterion for significance was established *a priori* at the 95% confidence level (p < 0.05).

## 3.0 Results

### 3.1 Heron vaccine effects on antinociceptive potency

Across all baseline sessions after vehicle administration and prior to drug administration, tail-withdrawal latencies were 19.9 ± 0.02 s and 4.9 ± 0.1 at 40°C and 50°C, respectively. Figure 1 shows active and control vaccine effects in cohorts of rats receiving either SC or IV opioid agonist administration. MOR ligand ED_50_ values during each treatment condition are reported in Tables 1 and 2. SC heroin antinociceptive potency was significantly attenuated compared to baseline at weeks 3, 6 and 8 (time: F_1.9, 9.4_=9.68, p=0.0056), whereas SC fentanyl antinociceptive potency was not significantly altered (Panel A). Similarly, IV heroin antinociceptive potency in vaccinated rats was significantly attenuated compared to baseline at weeks 3, 5 and 7 (time: F_1.2, 7.1_=13.65, p<0.05); however, heroin antinociceptive potency was not altered following control vaccine administration, and IV fentanyl antinociceptive potency was not significantly altered following active or control vaccine administration (Panel B). Maximum potency shifts regardless of time following active heroin vaccine administration in group 1 (SC) and group 2 (IV) were not significantly different (Panel C). Post-hoc power analyses indicated the experiments were underpowered (calculated power=0.42) to detect a significant difference between SC and IV groups. To achieve a power = 0.8 for this experiment, an additional 16 animals (8 per route of administration group) would need to be tested. Midpoint titers were similar over time in the SC and IV cohorts (Panel D) and correlated with antinociceptive potency shifts of SC heroin (F_1,16_ = 5.37, p=0.03; R^2^ = 0.25), but not IV heroin (Supplementary Figure 1). 6-AM antinociceptive potency was significantly attenuated at weeks 4 and 6 compared to baseline (time: F_2.1, 9.4_=10.72, p<0.05) (Panel E). For morphine, there was a main effect of time (F_2.2, 9.8_=6.4, p<0.05), but post-hoc tests failed to detect a significant difference at any time point (Panel F).

**Table 1.**
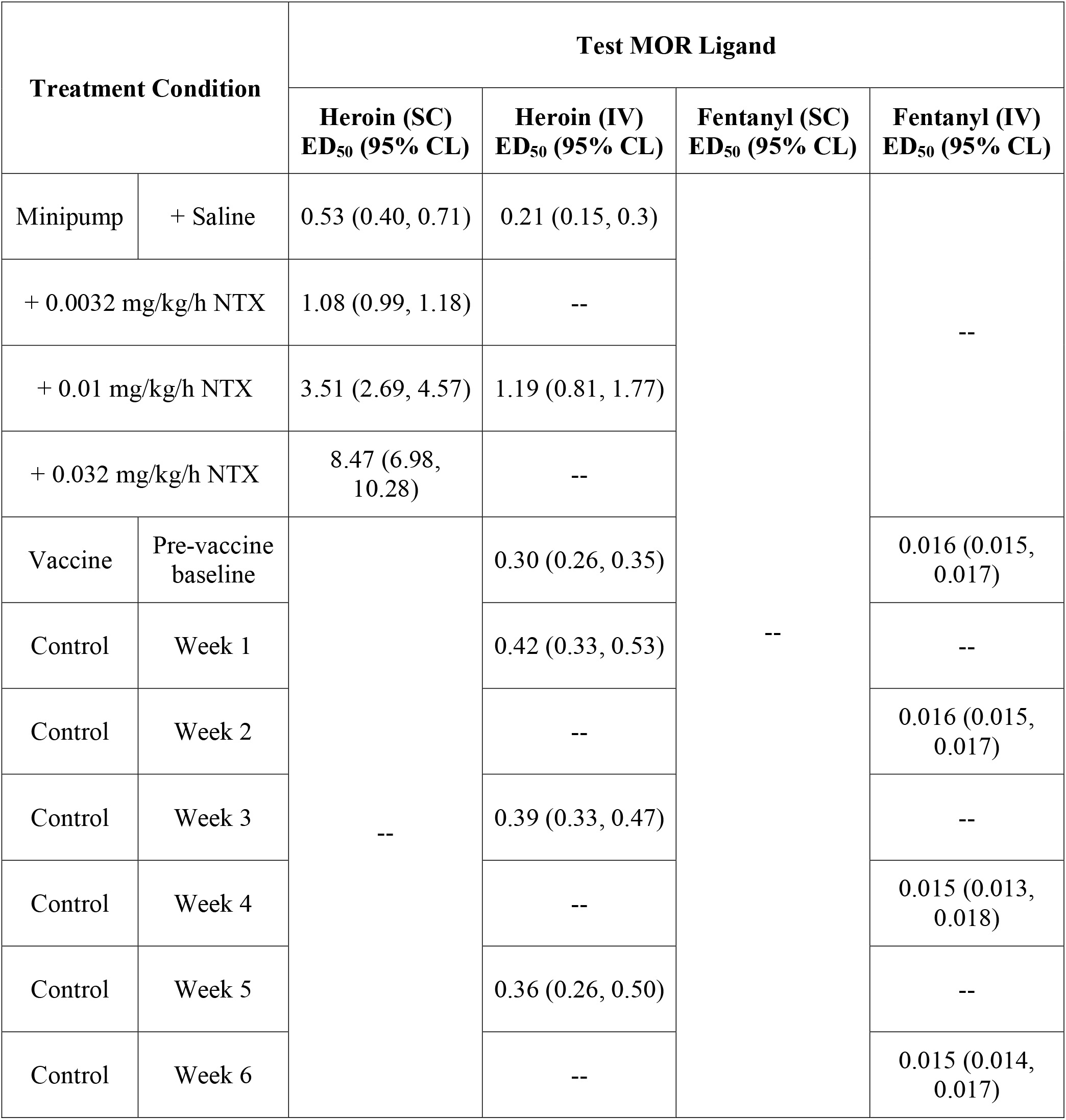
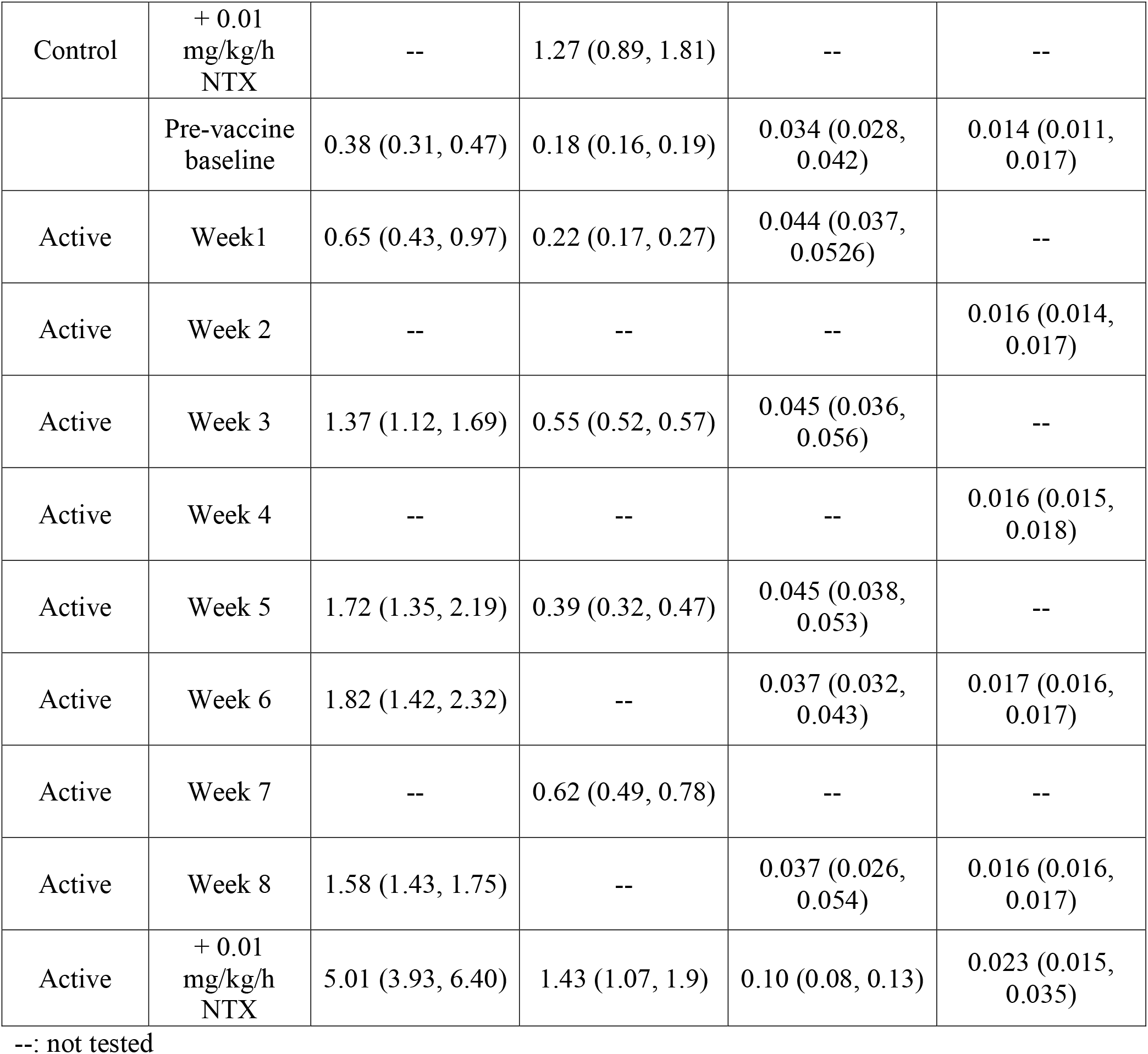
Group mean MOR ligand ED_50_ values and (95% confidence limits; CL) in the warm water tail withdrawal procedure during continuous naltrexone (NTX) or vaccine treatment (n=5-6 rats).

**Table 2.**
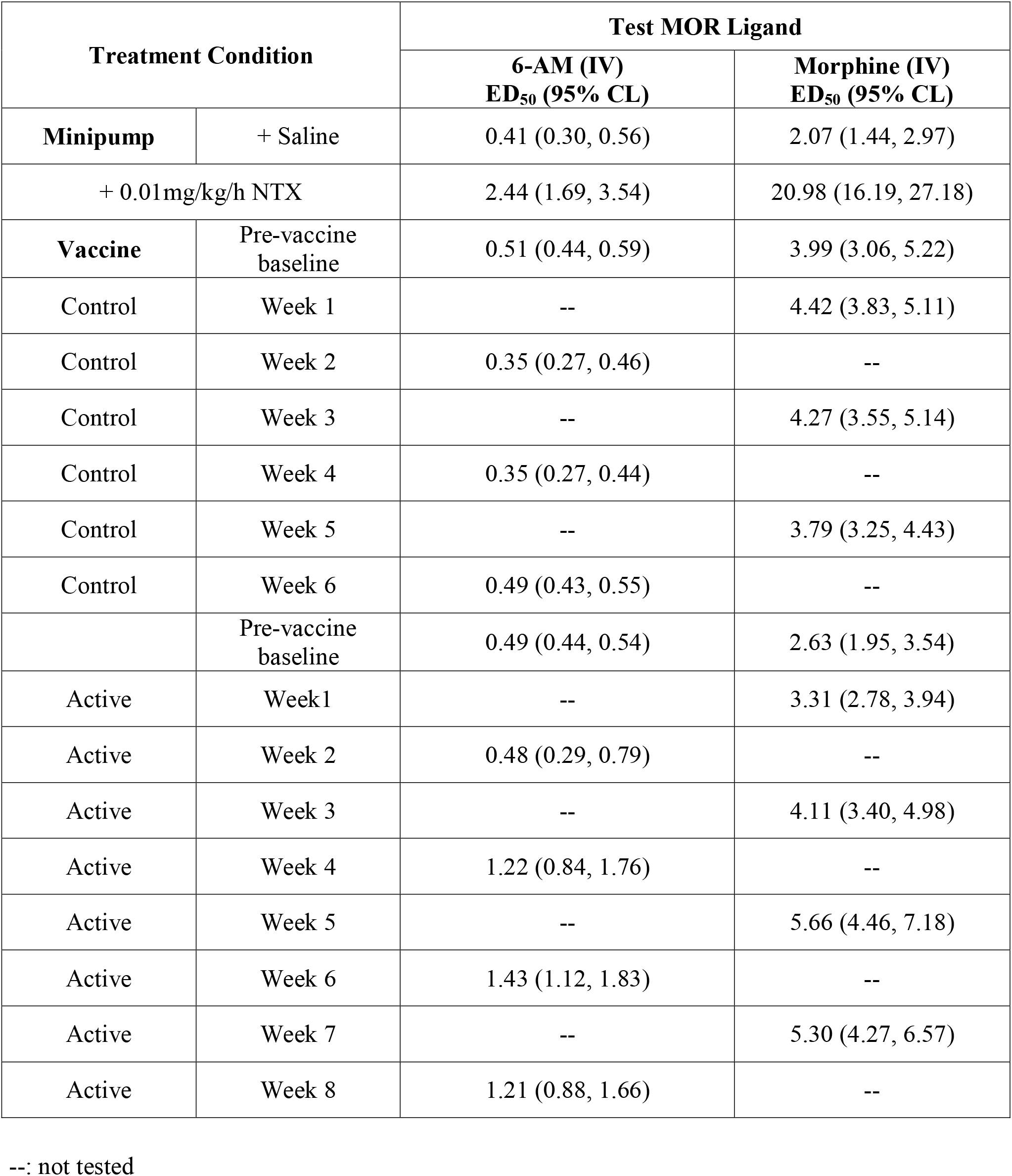
Group mean MOR ligand ED_50_ values and (95% confidence limits; CL) in the warm-water tail-withdrawal procedure during continuous naltrexone (NTX) or vaccine treatment (n=5-6 rats).

| Treatment Condition |  | Test MOR Ligand |  |
| --- | --- | --- | --- |
|  |  | 6-AM (IV)<br>ED<sub>50</sub> (95% CL) | Morphine (IV)<br>ED<sub>50</sub> (95% CL) |
| **Minipump** | + Saline | 0.41 (0.30, 0.56) | 2.07 (1.44, 2.97) |
| + 0.01mg/kg/h NTX |  | 2.44 (1.69, 3.54) | 20.98 (16.19, 27.18) |
| **Vaccine** | Pre-vaccine baseline | 0.51 (0.44, 0.59) | 3.99 (3.06, 5.22) |
| Control | Week 1 | -- | 4.42 (3.83, 5.11) |
| Control | Week 2 | 0.35 (0.27, 0.46) | -- |
| Control | Week 3 | -- | 4.27 (3.55, 5.14) |
| Control | Week 4 | 0.35 (0.27, 0.44) | -- |
| Control | Week 5 | -- | 3.79 (3.25, 4.43) |
| Control | Week 6 | 0.49 (0.43, 0.55) | -- |
|  | Pre-vaccine baseline | 0.49 (0.44, 0.54) | 2.63 (1.95, 3.54) |
| Active | Week1 | -- | 3.31 (2.78, 3.94) |
| Active | Week 2 | 0.48 (0.29, 0.79) | -- |
| Active | Week 3 | -- | 4.11 (3.40, 4.98) |
| Active | Week 4 | 1.22 (0.84, 1.76) | -- |
| Active | Week 5 | -- | 5.66 (4.46, 7.18) |
| Active | Week 6 | 1.43 (1.12, 1.83) | -- |
| Active | Week 7 | -- | 5.30 (4.27, 6.57) |
| Active | Week 8 | 1.21 (0.88, 1.66) | -- |

--: not tested

**Figure 1.**
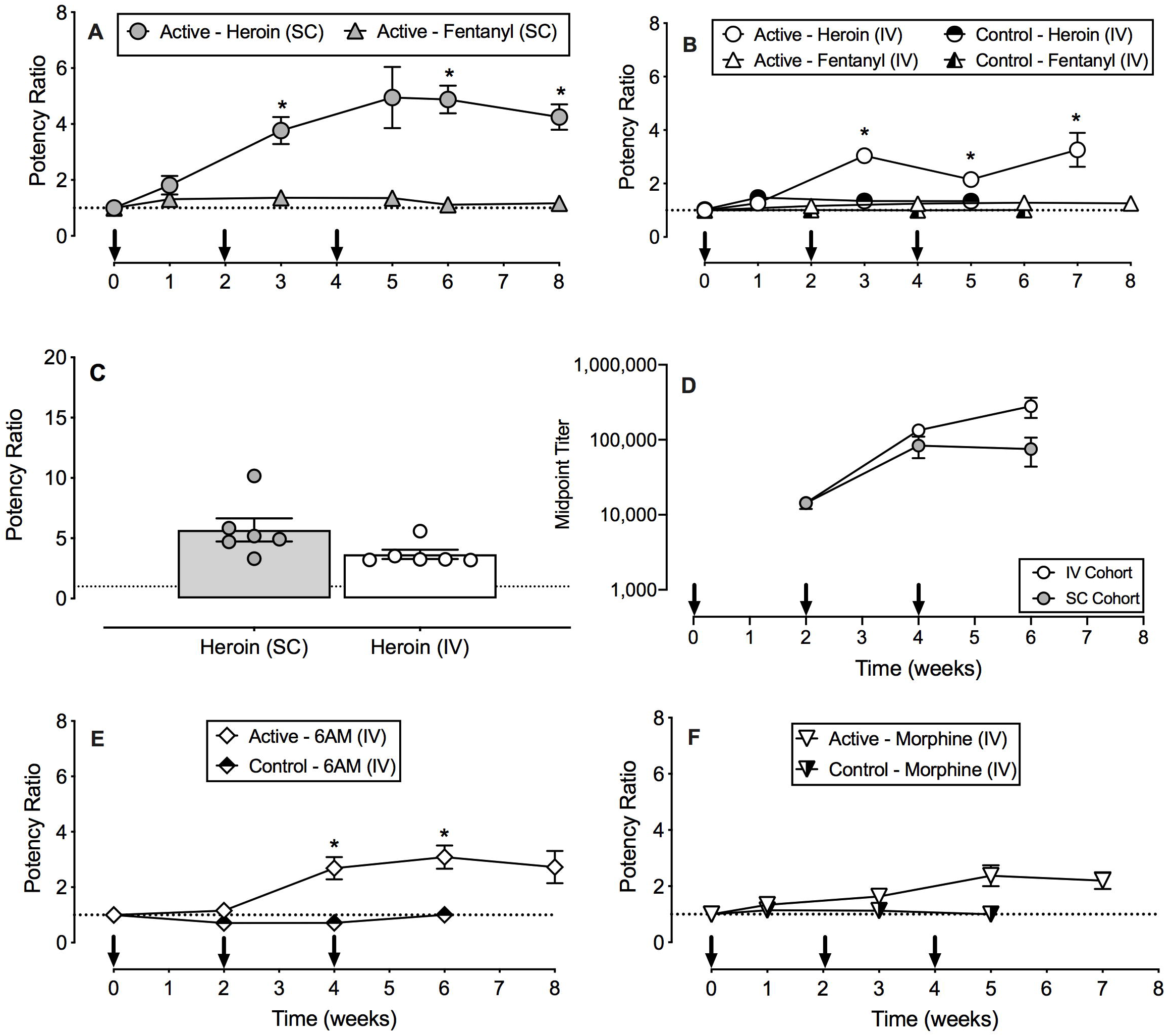
Effects of a heroin-tetanus toxoid (TT) conjugate vaccine on the antinociceptive potency of heroin, 6-acetylmorphine (6-AM), morphine, and fentanyl in male and female rats. Panels A and B show effects of active or control vaccine on subcutaneous (SC) and intravenous (IV) administered heroin and fentanyl antinociceptive potency. Panel C shows individual subject and group mean maximum active heroin vaccine effects between SC and IV administered heroin. Panel D shows midpoint titer levels as a function of time in both the IV and SC cohorts that received active vaccine. Panels E and F show effects of active and control vaccine on IV administered 6-AM, and morphine antinociceptive potency. Abscissae: time in weeks (Panels A, B, D, E, and F) and drug administered and route of administration (Panel C). Ordinate: potency ratio (Panels A, B, C, E, and F) and midpoint titer (Panel E). Arrows in Panels A, B, D, E, and F indicate when either active or control vaccine was administered. Asterisks indicate statistical significance (p<0.05) compared to week 0. All points in panels A, B, D, E, and F and bars in panel C represent mean ± s.e.m. of 5-6 rats.

### 3.2 Chronic naltrexone treatment effects on antinociceptive potency

For comparison to vaccine effects, Figure 2 shows chronic naltrexone treatment effects on opioid agonist dose-effect functions and potency ratios. Because there were no significant differences in the opioid agonist ED_50_ values between treatment days 7 and 14 for any naltrexone dose, data were averaged for both statistical analyses and graphical display. MOR ligand ED_50_ values during each naltrexone treatment condition are reported in Tables 1 and 2. Naltrexone (0.0032-0.032 mg/kg/h) treatment produced dose-dependent rightward shifts in cumulative SC heroin dose-effect functions (Panel A) and dose-dependent increases in SC heroin potency ratios (Panel B; (naltrexone dose: F_3,20_=76.36, p<0.0001)). Naltrexone plasma levels were 2.0 ± 0.5 ng/mL during 0.01 mg/kg/h naltrexone treatment and 4.7 ± 0.4 ng/mL during 0.032 mg/kg/h naltrexone treatment. Panels C, D, and E show that 0.01 mg/kg/h naltrexone treatment also produced rightward shifts in dose-effect curves for IV heroin, IV 6-AM, and IV morphine antinociception, respectively. Panel F shows that 0.01 mg/kg/h naltrexone produced similar potency ratios for SC heroin, IV heroin, IV 6-AM and IV morphine.

**Figure 2.**
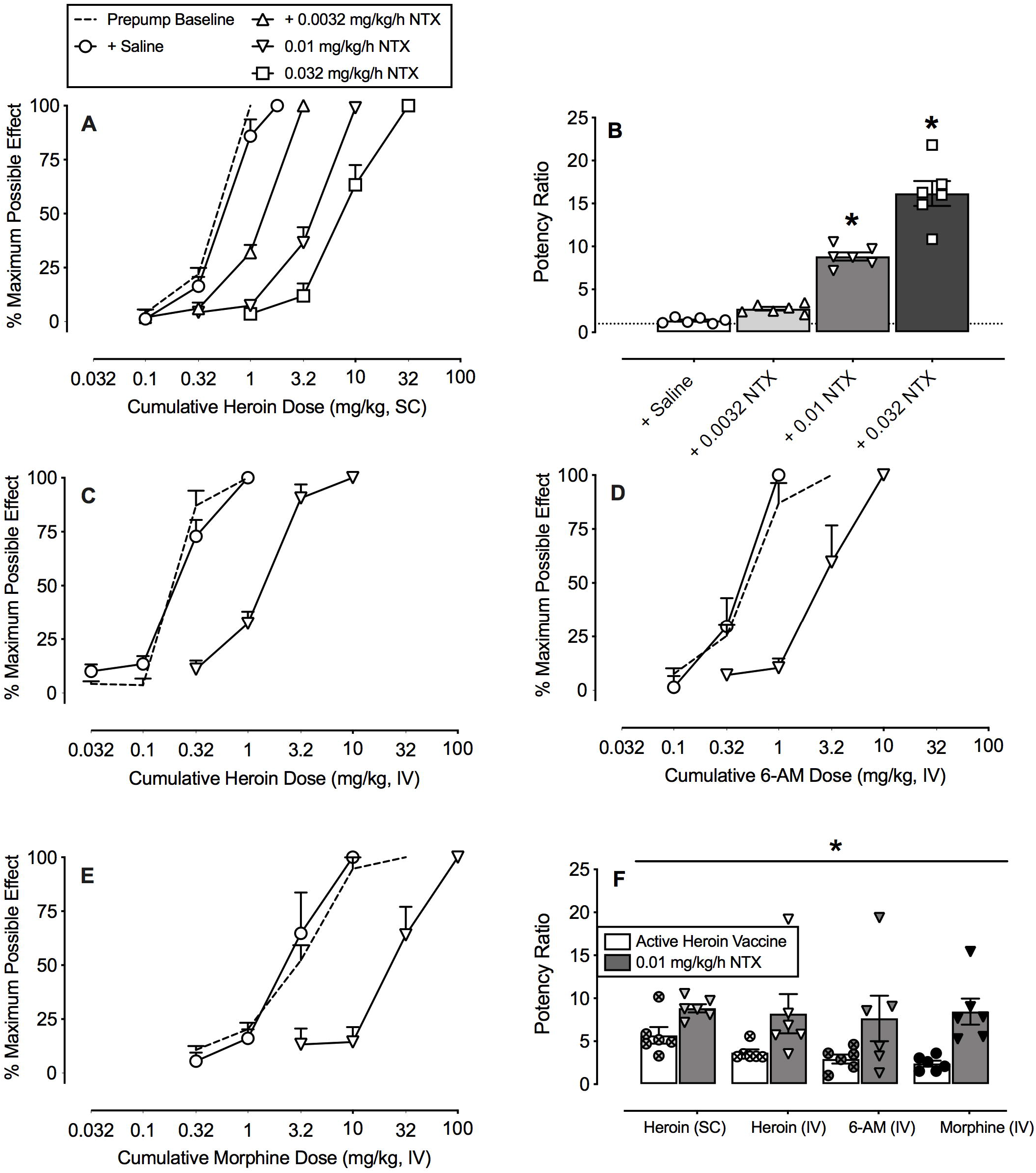
Effects of continuous naltrexone treatment on the antinociceptive effects of heroin, 6-acetylmorphine, morphine and fentanyl in male and female rats. Panel A shows effects of saline and naltrexone (NTX; 0.0032-0.032 mg/kg/h) osmotic pumps on the antinociceptive potency of subcutaneous (SC) heroin. Panel B shows individual subject and group mean potency ratios for SC heroin following saline and naltrexone treatments. Panel C, D and E shows effects of 0.01 mg/kg/h naltrexone osmotic pumps on the antinociceptive effects of intravenous (IV) heroin, 6-AM and morphine, respectively. Panel F shows individual subject and group mean maximum treatment effects for the active heroin vaccine and 0.01 mg/kg/h naltrexone treatments across different routes of heroin administration (SC or IV) and opioid agonists (heroin, 6-AM or morphine). Abscissae: cumulative drug dose in milligrams per kilogram, IV or SC (Panels A, C, D, and E), naltrexone treatment condition (Panel B), drug administered and route of administration (Panel F). Ordinate: Percent Maximum Possible Effect (Panels A, C, D, and E), Potency Ratio (Panels B and F). Asterisks indicate a statistical significance (p<0.05) compared to saline (Panel B), or a main effect of treatment (Panel F). All points in panels A, C, D, and E and bars in panels B and F represent mean ± s.e.m. of 5-6 rats.

Clinically available doses of depot naltrexone produce minimal plasma naltrexone levels of approximately 2 ng/ml and ≥ 8-fold potency shifts in heroin self-administration in humans (Bigelow et al., 2012; Comer et al., 2006; Sullivan et al., 2006). In the present study, these parameters for naltrexone plasma levels and heroin potency ratios were approximated by the 0.01 mg/kg/h naltrexone dose. Accordingly, Figure 2F also compares effects of 0.01 mg/kg/h naltrexone with maximal potency ratios achieved with active heroin vaccine. Because there was only a main effect of treatment condition (F_1,10_=42.8, p<0.0001) and no main effect of test drug or interaction, the statistical analysis defaulted to an unpaired t-test. The 0.01 mg/kg/h naltrexone dose produced greater potency ratios than the active heroin vaccine for all opioid agonists (t_3.7_=6.1, p=0.0048).

### 3.3 Naltrexone effectiveness in active heroin vaccinated rats

Figure 3 compares heroin potency ratios produced by 0.01 mg/kg/h naltrexone in unvaccinated rats and in Cohorts 1 (SC heroin: Panels A and B) and 2 (IV heroin: Panels C and D) active heroin vaccine-treated rats. The SC heroin potency ratio produced by 0.01 mg/kg/h naltrexone was significantly greater in the active heroin vaccine cohort compared to the unvaccinated rats (Panel B: t_6.0_=2.72, p=0.03). In contrast, the IV heroin potency ratios were not significantly different between active heroin vaccine and unvaccinated rats treated with 0.01 mg/kg/h naltrexone.

**Figure 3.**
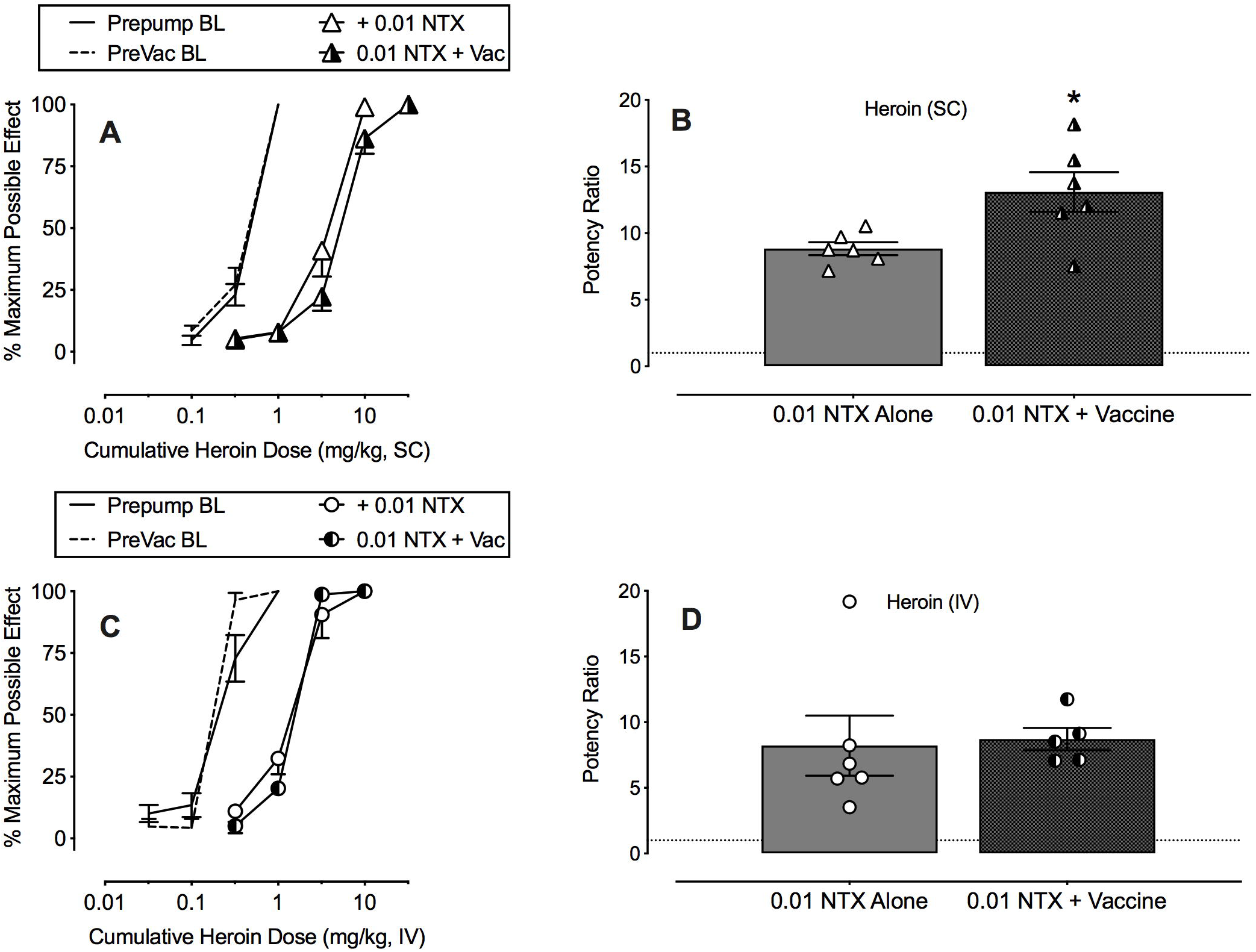
Effectiveness of 0.01 mg/kg/h naltrexone (NTX) alone or in combination with the active heroin vaccine on the antinociceptive potency of subcutaneous (SC) or intravenous (IV) heroin in male and female rats. Panel A shows cumulative SC heroin dose-effect functions at baseline (BL) and during 0.01 mg/kg/h naltrexone alone or 0.01 mg/kg/h naltrexone plus active vaccine (week 9). Panel B shows corresponding individual and group mean potency shifts compared to BL. Panel C shows cumulative IV heroin dose-effect functions at BL and during 0.01 mg/kg/h naltrexone alone or 0.01 mg/kg/h naltrexone plus active vaccine (week 11). Panel D shows corresponding individual and group mean potency shifts compared to BL. Abscissae: cumulative drug dose (mg/kg, SC or IV) (Panels A and C) or treatment condition (Panels B and D). Ordinate: percent maximum possible effect (Panels A and C) or potency ratios (Panels B and D). Asterisk indicates statistical significance (p<0.05) compare to naltrexone alone. All points in panels A and C and bars in panels B and D represent mean ± s.e.m. of 5-6 rats.

## 4.0 Discussion

The aim of the present study was to determine whether route of heroin administration impacted the effectiveness of a heroin-TT conjugate vaccine to attenuate heroin antinociceptive potency in male and female rats. In addition, heroin vaccine effectiveness was compared to continuous naltrexone treatment to model depot naltrexone formulations used clinically for OUD treatment. There were four main findings. First, the heroin vaccine was effective at blunting the antinociceptive effects of both SC and IV heroin and the heroin vaccine was selective for heroin and 6-AM vs. morphine and fentanyl. In contrast, the control vaccine did not alter the antinociceptive effects of heroin, 6-AM, morphine, or fentanyl. Second, heroin vaccine effectiveness as a consequence of heroin route of administration was unable to be fully determined as absent due to statistical power below 0.8. Third, naltrexone treatment decreased the antinociceptive potency of heroin, 6-AM and morphine similarly and the minimally effective naltrexone dose produced an approximate 8-fold potency shift and plasma naltrexone levels around 2 ng/mL. Furthermore, naltrexone treatment was more effective than this heroin vaccine formulation. Finally, the heroin vaccine enhanced naltrexone effectiveness to attenuate heroin antinociceptive potency, but only when heroin was administered SC, not IV. Taken together, these results suggest that the effectiveness of this heroin-vaccination regimen may not be sufficient to translate this heroin vaccine formulation as an OUD monotherapy.

Maximum antinociceptive potency shifts following vaccine administration were approximately 3- and 5-fold for IV and SC heroin, respectively and lasted for at least seven-to-eight weeks. The present results are consistent with previous preclinical vaccine studies in male and female mice (Bremer et al., 2017; Bremer et al., 2014; Hwang et al., 2018a; Hwang et al., 2018b; Hwang et al., 2018c; Jalah et al., 2015; Matyas et al., 2014; Sulima et al., 2017) and male rats (Raleigh et al., 2018; Raleigh et al., 2013; Schlosburg et al., 2013; Stowe et al., 2011; Sulima et al., 2017) that heroin-targeted vaccines attenuate heroin-related behavioral effects. The present results extended these previous findings to the inclusion of female rats and no sex difference in vaccine effectiveness was observed. In addition, midpoint antibody titer levels in the present study were comparable to previous levels in mice (Bremer et al., 2017). The magnitude of heroin vaccine-induced potency shifts was similar to previous heroin-TT conjugate vaccine results using schedule-controlled responding in nonhuman primates (4-fold) and antinociceptive effects in female mice (5-7 fold) for heroin administered intramuscular (IM) and IP, respectively (Bremer et al., 2017; Hwang et al., 2018b; Hwang et al., 2018c). However, heroin potency shifts observed in the present study following either SC or IV heroin vaccine administration were less than previously reported antinociceptive potency shifts (12-14 fold) in male mice administered heroin SC and IP (Bremer et al., 2017). One potential reason for the variability in heroin vaccine effectiveness may be due to vaccine formulation differences such as hapten copy number or adjuvant and conjugate dosing (Bremer et al., 2017; Hwang et al., 2018a).

The present study utilized opioid-induced antinociception as a surrogate endpoint to characterize heroin vaccine effectiveness and selectivity in blocking heroin effects. Ultimately, of course, the goal for use of any vaccine would not be to block the antinociceptive or analgesic effects of an opioid agonist, but rather to block abuse-related or toxic effects of opioid agonists. However, preclinical antinociceptive procedures, such as warm-water tail-withdrawal, have been utilized for decades to efficiently characterize in vivo potency, time course, and receptor mechanisms of opioid compounds (Whiteside et al., 2013, 2016). A recent study examining fentanyl vaccine effectiveness reported similar vaccine potency shifts for fentanyl self-administration under a fentanyl vs. food choice procedure and fentanyl antinociception in the same rats (Townsend et al., 2019a). These results support the utility of opioid antinociception as an efficient surrogate dependent measure of vaccine effectiveness to block opioid agonist effects in general. Further research that evaluates vaccine effectiveness across multiple dependent measures, including abuse-related effects and respiratory depression, is clearly warranted to better characterize the clinical potential and utility of opioid vaccines.

In addition to considering the utility of antinociception as a useful surrogate endpoint for vaccine evaluation, translational research should also consider the utility of rats as model organisms to predict vaccine effectiveness in humans. In particular, the heroin vaccine studied here has different affinities for heroin, 6AM, and morphine, and translation of vaccine effectiveness to block heroin effects may be influenced by species differences in the role of heroin, 6AM, and morphine in mediating heroin effects. Some data suggest that morphine may play an especially prominent role in mediating heroin effects in humans (Inturrisi et al., 1984; Rook et al., 2006). At present, it is unclear if morphine is more important as an active heroin metabolite in humans than in rats. However, if this were the case, and given the relatively low affinity of the present vaccine for morphine, it is possible that this vaccine may have even lower effectiveness to block heroin effects in humans than in rats.

Heroin vaccine effectiveness to blunt heroin antinociceptive potency was not significantly different between the SC and IV routes of administration; however, a post-hoc power analyses indicated that the experiment was not sufficiently powered to conclude that the absence of a significant difference between heroin route of administration was not a false negative. A previous study suggested route of administration (SC vs. IV) was an important factor for oxycodone and heroin vaccine effectiveness in rats (Raleigh et al., 2018). One key difference between the present study and Raleigh (2018) is the heroin metabolite that was targeted by the vaccine. The heroin vaccine tested in the present study produced antibodies that have the highest affinity for 6-AM (Bremer et al., 2017) because 6-AM is hypothesized to be the primary mediator of heroin pharmacodynamics in mice (Andersen et al., 2009), rats (Gottås et al., 2014; Inturrisi et al., 1983), and humans (Inturrisi et al., 1984; Rook et al., 2006). In contrast, the heroin vaccine in Raleigh (2018) targeted the heroin metabolite morphine and displayed cross-reactivity with 6-AM (Raleigh et al., 2013). Heroin potency shifts were not examined in the Raleigh (2018) study preventing a direct comparison of vaccine effectiveness between studies. Ultimately, human laboratory studies examining heroin vaccine effectiveness will provide critical feedback regarding which heroin metabolite to target with an immunopharmacotherapy.

Selectivity for the target abused opioid is one potential advantage of immunopharmacotherapies over current FDA-approved treatments buprenorphine and naltrexone because the immunopharmacotherapy would allow clinicians to use structurally dissimilar MOR agonists for pain management (Banks et al., 2018; Bremer et al., 2017). However, high selectivity may also be a disadvantage of immunopharmacotherapies because a motivated individual may switch to abusing a structurally dissimilar MOR agonist not targeted by the vaccine. In addition, heroin has been reported to be contaminated with fentanyl or fentanyl analogs which would also mitigate vaccine effectiveness (Hibbs et al., 1991; Stogner, 2014). As a result, recent preclinical studies have examined the effectiveness of a combination heroin/fentanyl vaccine (Hwang et al., 2018b; Hwang et al., 2018c).

Continuous naltrexone treatment dose-dependently antagonized the antinociceptive effects of heroin. Naltrexone treatment produced similar antinociceptive potency shifts for heroin and its metabolites 6-AM and morphine. These results were consistent with previous studies showing chronic naltrexone antagonized other behavioral effects of opioid agonists (Dean et al., 2008; Sakloth and Negus, 2018). Naltrexone plasma levels were dose-dependent in the present study, but maximum naltrexone levels were less than levels after naltrexone extended-release (XR-NTX) formulation administration in rats (Todtenkopf et al., 2009). Although higher naltrexone plasma levels were achieved with the XR-NTX formulation, clinical studies have suggested that clinical effectiveness was achieved with a minimal naltrexone level of 2 ng/ml (Bigelow et al., 2012; Comer et al., 2006). Consistent with these human studies, a dose of 0.01 mg/kg/h naltrexone produced this minimal naltrexone plasma level and an approximate 8-fold antinociceptive potency shift for SC and IV heroin, IV 6-AM, and IV morphine.

Regardless of heroin route of administration, the heroin-TT conjugate vaccine formulation evaluated in the present study was less effective than the minimally effective continuous naltrexone dose. These results suggest limited clinical effectiveness of this specific heroin-TT conjugate vaccine as an OUD monotherapy utilizing the current adjuvant, immune modulators, and conjugate dosing regimen. However, the present results do support continued research development and optimization of a heroin vaccine to enhance vaccine effectiveness. Furthermore, immunopharmacotherapies could also be clinically utilized as adjunctive therapies to current FDA-approved OUD treatments to provide enhanced protection against opioid overdose or potentially increase dosing intervals for extended release naltrexone or buprenorphine. For example, a nicotine vaccine was co-administered with the nicotinic partial agonist varenicline in a randomized placebo-controlled clinical trial (Hoogsteder et al., 2014). Compared to the placebo vaccine plus varenicline group, the nicotine vaccine plus varenicline was not more effective compared to the placebo vaccine plus varenicline group, similar to the present heroin vaccine results when heroin was administered IV. Overall, the present results do provide preliminary empirical evidence that combining immunopharmacotherapies with current FDA-approved OUD pharmacotherapies warrants further evaluation to address the ongoing opioid crisis.

## References

Andersen, J.M., Ripel, Å., Boix, F., Normann, P.T., Mørland, J., 2009. Increased Locomotor Activity Induced by Heroin in Mice: Pharmacokinetic Demonstration of Heroin Acting as a Prodrug for the Mediator 6-Monoacetylmorphine in Vivo. J Pharmacol Exp Ther 331, 153–161.

Anton, B., Leff, P., 2006. A novel bivalent morphine/heroin vaccine that prevents relapse to heroin addiction in rodents. Vaccine 24, 3232–3240.

Baehr, C., Pravetoni, M., 2019. Vaccines to treat opioid use disorders and to reduce opioid overdoses. Neuropsychopharmacology 44, 217.

Banks, M.L., Olson, M.E., Janda, K.D., 2018. Immunopharmacotherapies for Treating Opioid Use Disorder. Trends Pharmacol Sci 39, 908–911.

Barbosa-Leiker, C., McPherson, S., Layton, M.E., Burduli, E., Roll, J.M., Ling, W., 2018. Sex differences in opioid use and medical issues during buprenorphine/naloxone treatment. Am J Drug Alcohol Abuse 44, 488–496.

Becker, J.B., Koob, G.F., 2016. Sex Differences in Animal Models: Focus on Addiction. Pharmacol Rev 68, 242–263.

Bigelow, G.E., Preston, K.L., Schmittner, J., Dong, Q., Gastfriend, D.R., 2012. Opioid challenge evaluation of blockade by extended-release naltrexone in opioid-abusing adults: Dose– effects and time-course. Drug Alcohol Depend 123, 57–65.

Bonese, K.F., Wainer, B.H., Fitch, F.W., Rothberg, R.M., Schuster, C.R., 1974. Changes in heroin self-administration by a rhesus monkey after morphine immunisation. Nature 252, 708–710.

Bremer, P.T., Schlosburg, J.E., Banks, M.L., Steele, F.F., Zhou, B., Poklis, J.L., Janda, K.D., 2017. Development of a Clinically Viable Heroin Vaccine. J Am Chem Soc 139, 8601–8611.

Bremer, P.T., Schlosburg, J.E., Lively, J.M., Janda, K.D., 2014. Injection route and TLR9 agonist addition significantly impact heroin vaccine efficacy. Mol Pharm 11, 1075–1080.

Comer, S.D., Sullivan, M.A., Yu, E., Rothenberg, J.L., Kleber, H.D., Kampman, K., Dackis, C., O’Brien, C.P., 2006. Injectable, sustained-release naltrexone for the treatment of opioid dependence: a randomized, placebo-controlled trial. Arch Gen Psych 63(2), 210–218.

Council, N.R., 2011. Guide for the care and use of laboratory animals, 8th ed. National Academies Press, Washington DC.

Dean, R.L., Todtenkopf, M.S., Deaver, D.R., Arastu, M.F., Dong, N., Reitano, K., O’driscoll, K., Kriksciukaite, K., Gastfriend, D.R., 2008. Overriding the blockade of antinociceptive actions of opioids in rats treated with extended-release naltrexone. Pharmacol Biochem Behav 89(4), 515–522.

Gottås, A., Boix, F., Øiestad, E.L., Vindenes, V., Mørland, J., 2014. Role of 6-monoacetylmorphine in the acute release of striatal dopamine induced by intravenous heroin. Int J Neuropsychopharmacol 17, 1357–1365.

Gottås, A., Øiestad, E., Boix, F., Vindenes, V., Ripel, Å., Thaulow, C., Mørland, J., 2013. Levels of heroin and its metabolites in blood and brain extracellular fluid after iv heroin administration to freely moving rats. Br J Pharmacol 170, 546–556.

Hedegaard, H., Warner, M., Miniño, A.M., 2018. Drug overdose deaths in the United States, 1999-2017. NCHS Data Brief 329, 1–8.

Hibbs, J., Perper, J., Winek, C.L.J.J., 1991. AN outbreak of designer drug—related deaths in pennsylvania. JAMA 265, 1011–1013.

Hoogsteder, P.H., Kotz, D., van Spiegel, P.I., Viechtbauer, W., van Schayck, O.C., 2014. Efficacy of the nicotine vaccine 3′‐AmNic‐rEPA (NicVAX) co‐administered with varenicline and counselling for smoking cessation: a randomized placebo‐controlled trial. Addiction 109, 1252–1259.

Hwang, C.S., Bremer, P.T., Wenthur, C.J., Ho, S.O., Chiang, S., Ellis, B., Zhou, B., Fujii, G., Janda, K.D., 2018a. Enhancing Efficacy and Stability of an Antiheroin Vaccine: Examination of Antinociception, Opioid Binding Profile, and Lethality. Mol Pharm 15, 1062–1072.

Hwang, C.S., Smith, L.C., Natori, Y., Ellis, B., Zhou, B., Janda, K.D., 2018b. Efficacious vaccine against heroin contaminated with fentanyl. ACS Chem Neurosci 9, 1269–1275.

Hwang, C.S., Smith, L.C., Natori, Y., Ellis, B., Zhou, B., Janda, K.D., 2018c. Improved Admixture Vaccine of Fentanyl and Heroin Hapten Immunoconjugates: Antinociceptive Evaluation of Fentanyl-Contaminated Heroin. ACS Omega 3, 11537–11543.

Inturrisi, C.E., Max, M.B., Foley, K.M., Schultz, M., Shin, S.-U., Houde, R.W., 1984. The Pharmacokinetics of Heroin in Patients with Chronic Pain. N Engl J Med 310, 1213–1217.

Inturrisi, C.E., Schultz, M., Shin, S., Umans, J.G., Angel, L., Simon, E.J., 1983. Evidence from opiate binding studies that heroin acts through its metabolites. Life Sci 33, 773–776.

Jaffe, J.H., O’Keeffe, C., 2003. From morphine clinics to buprenorphine: regulating opioid agonist treatment of addiction in the United States. Drug Alcohol Depend 70, S3–S11.

Jalah, R., Torres, O.B., Mayorov, A.V., Li, F., Antoline, J.F., Jacobson, A.E., Rice, K.C., Deschamps, J.R., Beck, Z., Alving, C.R., 2015. Efficacy, but not antibody titer or affinity, of a heroin hapten conjugate vaccine correlates with increasing hapten densities on tetanus toxoid, but not on CRM197 carriers. Bioconj Chem 26, 1041–1053.

Jarvis, B.P., Holtyn, A.F., De Fulio, A., Koffarnus, M.N., Leoutsakos, J.-M.S., Umbricht, A., Fingerhood, M., Bigelow, G.E., Silverman, K., 2019. The Effects of Extended-Release Injectable Naltrexone and Incentives for Opiate Abstinence in Heroin-Dependent Adults in a Model Therapeutic Workplace: A Randomized Trial. Drug Alcohol Depend 197, 220–227.

Jarvis, B.P., Holtyn, A.F., Subramaniam, S., Tompkins, D.A., Oga, E.A., Bigelow, G.E., Silverman, K., 2018. Extended‐release injectable naltrexone for opioid use disorder: a systematic review. Addiction 113, 1188–1209.

Lee, J.D., Nunes Jr, E.V., Novo, P., Bachrach, K., Bailey, G.L., Bhatt, S., Farkas, S., Fishman, M., Gauthier, P., Hodgkins, C.C., 2018. Comparative effectiveness of extended-release naltrexone versus buprenorphine-naloxone for opioid relapse prevention (X: BOT): a multicentre, open-label, randomised controlled trial. Lancet 391, 309–318.

Matyas, G.R., Rice, K.C., Cheng, K., Li, F., Antoline, J.F., Iyer, M.R., Jacobson, A.E., Mayorov, A.V., Beck, Z., Torres, O.B., 2014. Facial recognition of heroin vaccine opiates: type 1 cross-reactivities of antibodies induced by hydrolytically stable haptenic surrogates of heroin, 6-acetylmorphine, and morphine. Vaccine 32, 1473–1479.

Monico, L.B., Mitchell, S.G., 2018. Patient perspectives of transitioning from prescription opioids to heroin and the role of route of administration. Subst Abuse Treat Prevent Policy 13(1), 4.

Raleigh, M.D., Laudenbach, M., Baruffaldi, F., Peterson, S.J., Roslawski, M.J., Birnbaum, A.K., Carroll, F.I., Runyon, S.P., Winston, S., Pentel, P.R., 2018. Opioid dose-and route-dependent efficacy of oxycodone and heroin vaccines in rats. J Pharmacol Exp Ther 365, 346–353.

Raleigh, M.D., Pravetoni, M., Harris, A.C., Birnbaum, A.K., Pentel, P.R., 2013. Selective Effects of a Morphine Conjugate Vaccine on Heroin and Metabolite Distribution and Heroin-Induced Behaviors in Rats. J Pharmacol Exp Ther 344, 397–406.

Rook, E.J., Van Ree, J.M., Van Den Brink, W., Hillebrand, M.J.X., Huitema, A.D.R., Hendriks, V.M., Beijnen, J.H., 2006. Pharmacokinetics and Pharmacodynamics of High Doses of Pharmaceutically Prepared Heroin, by Intravenous or by Inhalation Route in Opioid-Dependent Patients. Basic Clin Pharmacol Toxicol 98, 86–96.

Sakloth, F., Negus, S.S., 2018. Naltrexone maintenance fails to alter amphetamine effects on intracranial self-stimulation in rats. Exp Clin Psychopharmacol 26, 195–204.

Saloner, B., Karthikeyan, S., 2015. Changes in substance abuse treatment use among individuals with opioid use disorders in the United States, 2004–2013. JAMA 314, 1515–1517.

SAMHSA, 2018. Key substance use and mental health indicators in the United States: Results from the National Survey on Drug Use and Health (HHS Publication No. SMA 18-5068, NSDUH Series H-53). Center for Behavioral Health Statistics and Quality, Substance Abuse and Mental Health Services Administration Rockville, MD.

Schlosburg, J.E., Vendruscolo, L.F., Bremer, P.T., Lockner, J.W., Wade, C.L., Nunes, A.A., Stowe, G.N., Edwards, S., Janda, K.D., Koob, G.F., 2013. Dynamic vaccine blocks relapse to compulsive intake of heroin. Proc Natl Acad Sci USA 110, 9036–9041.

Stogner, J.M., 2014. The potential threat of acetyl fentanyl: legal issues, contaminated heroin, and acetyl fentanyl “disguised” as other opioids. Ann Emerg Med 64, 637–639.

Stowe, G.N., Vendruscolo, L.F., Edwards, S., Schlosburg, J.E., Misra, K.K., Schulteis, G., Mayorov, A.V., Zakhari, J.S., Koob, G.F., Janda, K.D., 2011. A vaccine strategy that induces protective immunity against heroin. J Med Chem 54, 5195–5204.

Sulima, A., Jalah, R., Antoline, J.F., Torres, O.B., Imler, G.H., Deschamps, J.R., Beck, Z., Alving, C.R., Jacobson, A.E., Rice, K.C., 2017. A stable heroin analogue that can serve as a vaccine Hapten to induce antibodies that block the effects of heroin and its metabolites in rodents and that cross-react immunologically with related drugs of abuse. J Med Chem 61, 329–343.

Sullivan, M.A., Vosburg, S.K., Comer, S.D., 2006. Depot naltrexone: antagonism of the reinforcing, subjective, and physiological effects of heroin. Psychopharmacology 189, 37–46.

Tkacz, J., Severt, J., Cacciola, J., Ruetsch, C., 2012. Compliance with buprenorphine medication‐assisted treatment and relapse to opioid use. Am J Addict 21, 55–62.

Todtenkopf, M.S., O’Neill, K.S., Kriksciukaite, K., Turncliff, R.Z., Dean, R.L., Ostrovsky‐Day, I., Deaver, D.R., 2009. Route of administration affects the ability of naltrexone to reduce amphetamine‐potentiated brain stimulation reward in rats. Addict Biol 14, 408–418.

Townsend, E.A., Blake, S., Faunce, K.E., Hwang, C.S., Natori, Y., Zhou, B., Bremer, P.T., Janda, K.D., Banks, M.L., 2019a. Conjugate vaccine produces long-lasting attenuation of fentanyl vs. food choice and blocks expression of opioid withdrawal-induced increases in fentanyl choice in rats. Neuropsychopharmacology. doi: 10.1038/s41386-019-0385-9.

Townsend, E.A., Negus, S.S., Caine, S.B., Thomsen, M., Banks, M.L., 2019b. Sex differences in opioid reinforcement under a fentanyl vs. food choice procedure in rats. Neuropsychopharmacology. doi: 10.1038/s41386-019-0356-1.

Volkow, N.D., Collins, F.S., 2017. The role of science in addressing the opioid crisis. N Engl J Med 377, 391–394.

Whiteside, G.T., Pomonis, J.D., Kennedy, J.D., 2013. An industry perspective on the role and utility of animal models of pain in drug discovery. Neurosci Lett 557, 65–72.

Whiteside, G.T., Pomonis, J.D., Kennedy, J.D., 2016. Chapter Eleven - Preclinical Pharmacological Approaches in Drug Discovery for Chronic Pain, in: Barrett, J.E. (Ed.) Advances in Pharmacology. Academic Press, pp. 303–323.

