## Supplemental Figure 1 for "Effectiveness and selectivity of a heroin conjugate vaccine to attenuate heroin, 6-acetylmorphine, and morphine antinociception in rats: Comparison with naltrexone" (from another version of this preprint: version 1, 2019-03-15)

**Number of Figures:** 1

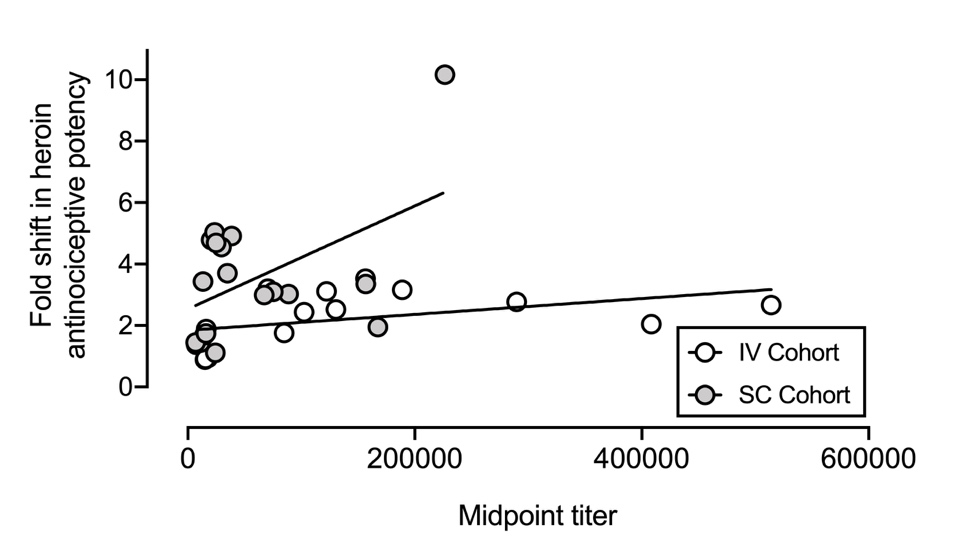

**Supplementary Figure 1.**

Individual maximum fold shift in heroin antinociceptive potency as a function of midpoint titer. Abscissae: Midpoint titer. Ordinate: Fold shift in antinociceptive potency.
